# Estimating parasite load dynamics to reveal novel resistance mechanisms to human malaria

**DOI:** 10.1101/321463

**Authors:** Michael T. Bretscher, Athina Georgiadou, Hyun Jae Lee, Michael Walther, Anna E. van Beek, Fadlila Fitriani, Diana Wouters, Taco W. Kuijpers, Davis Nwakanma, Umberto D’Alessandro, Eleanor M. Riley, Michael Levin, Lachlan J. Coin, Azra Ghani, David J. Conway, Aubrey J. Cunnington

**Affiliations:** Medical Research Council Centre for Outbreak Analysis and Modelling, Imperial College, London, United Kingdom; Section of Paediatrics, Imperial College, London, United Kingdom; Institute for Molecular Bioscience, University of Queensland, Brisbane, Australia; MRC Unit The Gambia at the London School of Hygiene and Tropical Medicine, Fajara, The Gambia; Department of Immunopathology, Sanquin Research and Landsteiner Laboratory of the Academic Medical Centre, University of Amsterdam, Amsterdam, the Netherlands; Department of Pediatric Hematology, Immunology and Infectious Diseases, Emma Children’s Hospital, Academic Medical Centre, Amsterdam, the Netherlands; Department of Blood Cell Research, Sanquin Research and Landsteiner Laboratory of the Academic Medical Centre, University of Amsterdam, Amsterdam, the Netherlands; The Roslin Institute and the Royal (Dick) School of Veterinary Studies, University of Edinburgh, Edinburgh, United Kingdom; Department of Immunology and Infection, London School of Hygiene and Tropical Medicine, London, United Kingdom; Department of Pathogen Molecular Biology, London School of Hygiene and Tropical Medicine, United Kingdom

**Author notes:** Corresponding author (AJC). Denotes equal contribution (Michael T. Bretscher, Athina Georgiadou, Hyun Jae Lee). Current address: Untere Grabenstraße 10, 88299 Leutkirch, Germany (Michael Walther). MTB performed the work related to this publication at Imperial College and is now an employee of F. Hoffmann-La Roche Ltd.

## Abstract

Improved methods are needed to identify host mechanisms which directly protect against human infectious diseases in order to develop better vaccines and therapeutics^1,2^. Pathogen load determines the outcome of many infections^3^, and is a consequence of pathogen multiplication rate, duration of the infection, and inhibition or killing of pathogen by the host (resistance). If these determinants of pathogen load could be quantified then their mechanistic correlates might be determined. In humans the timing of infection is rarely known and treatment cannot usually be withheld to monitor serial changes in pathogen load and host response. Here we present an approach to overcome this and identify potential mechanisms of resistance which control parasite load in *Plasmodium falciparum* malaria. Using a mathematical model of longitudinal infection dynamics for orientation, we made individualized estimates of parasite multiplication and growth inhibition in Gambian children at presentation with acute malaria and used whole blood RNA-sequencing to identify their correlates. We identified novel roles for secreted proteases cathepsin G and matrix metallopeptidase 9 (MMP9) as direct effector molecules which inhibit *P. falciparum* growth. Cathepsin G acts on the erythrocyte membrane, cleaving surface receptors required for parasite invasion, whilst MMP9 acts on the parasite. In contrast, the type 1 interferon response and expression of *CXCL10* (IFN-γ-inducible protein of 10 kDa, IP-10) were detrimental to control of parasite growth. Natural variation in iron status and plasma levels of complement factor H were determinants of parasite multiplication rate. Our findings demonstrate the importance of accounting for the dynamic interaction between host and pathogen when seeking to identify correlates of protection, and reveal novel mechanisms controlling parasite growth in humans. This approach could be extended to identify additional mechanistic correlates of natural- and vaccine-induced immunity to malaria and other infections.

## Main Text

To heuristically estimate the hidden dynamics of parasite load (Fig 1a) in naturally-infected malaria patients we calibrated a statistical prediction model using outputs from a mechanistic simulation which combined information from two datasets (Fig 1b). A historical dataset of the longitudinal course of infection in patients who were deliberately inoculated with *P. falciparum* as a treatment for neurosyphilis (malariatherapy) (Extended Data Figure 1) was used as a reference for changes in parasite load over time^4^. In this dataset, relationships were defined between measured and latent variables (which we assume to determine changes in parasite load over time), broadly based on a mathematical model proposed by Dietz et al.^4^. Measurements from Gambian children with malaria at the time of clinical presentation (Extended Data Table 1) were then used to estimate the values of the latent variables and dynamics of parasite load in individual children.

**Fig 1.**
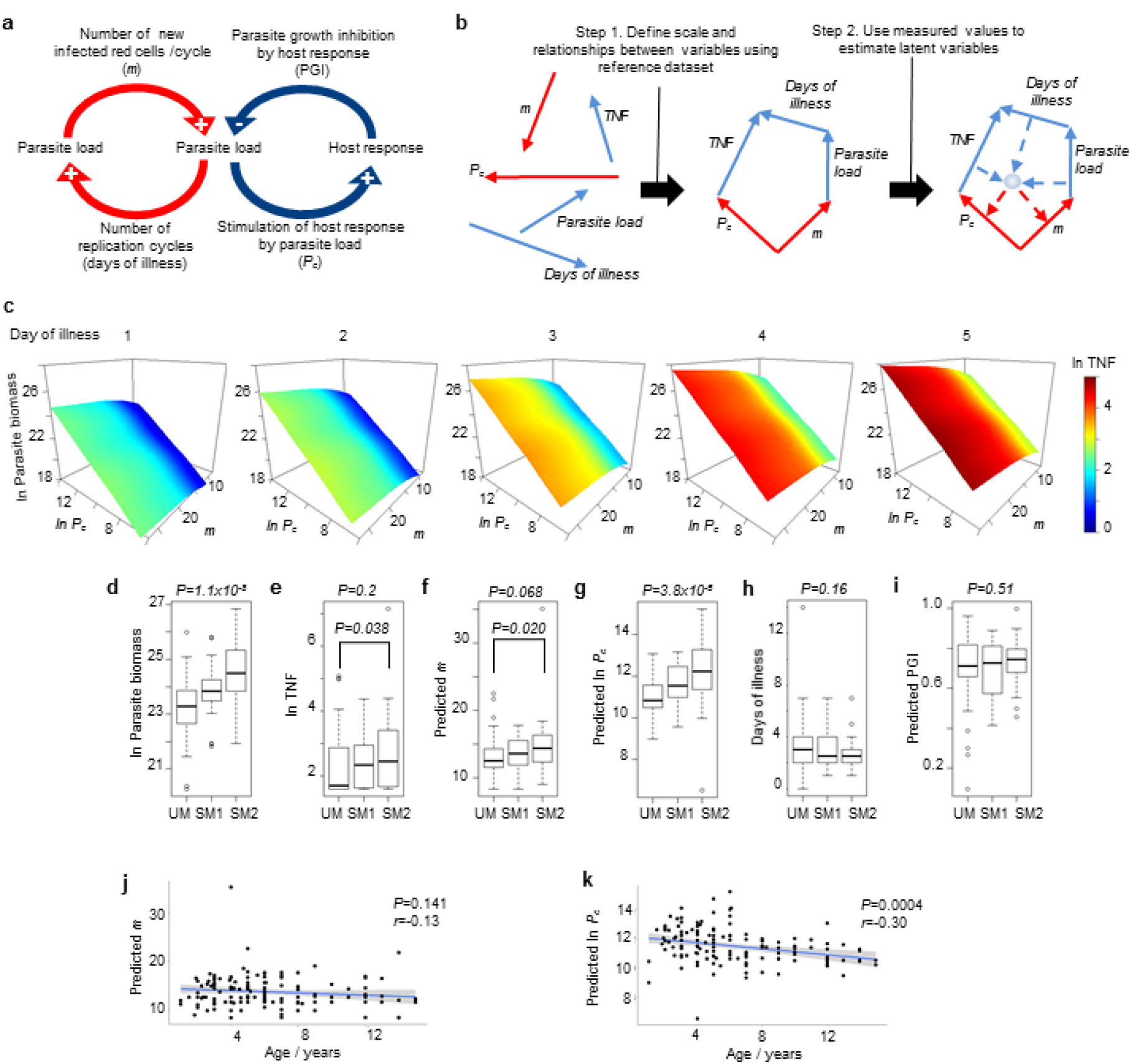
Estimating the dynamics of parasite load and host response in malaria. (**a, b**) Conceptual models of the major determinants of parasite load (**a**) and a framework for use of measurable variables at a single point in time (parasite load, duration of illness, plasma TNF, blue lines) to estimate latent variables (*m* and *P_c_*, red lines) (**b**). (**c**) Simulated relationships between *P_c_, m*, parasite biomass, duration of illness and TNF concentrations in Gambian children. (**d-i**) Comparisons of parasite biomass (**d**), TNF (**e**), predicted *m* (**f**), predicted *P_c_* (**g**), duration of illness (**h**), and predicted parasite growth inhibition (PGI, **i**), in 139 Gambian children with uncomplicated (UM, *n* =64) or severe malaria (SM1, prostration, *n* =36; SM2, any combination of cerebral malaria, hyperlactatemia or severe anemia, *n* =39). Boxes show median and interquartile range, whiskers extend 1.5-times the interquartile range or to limit of range; *P* for ANOVA (above plots), and for Mann-Witney test (UM vs SM2, within plots). (**j, k**) Correlation of predicted *m* (**j**) or *P_c_* (**k**) with age (blue line, linear regression; shaded area 95% confidence interval), *P* for Pearson correlation.

In our model, ascending parasite load dynamics (up to the first peak) in the malariatherapy reference data can be largely described with two individual-specific latent variables (Fig 1a, see Methods): the within-host multiplication rate, *m*, which is the initial rate of increase in parasite load before any constraint by the host response; and *P_c_*, which is defined by the parasite load required to stimulate a host response that reduces parasite growth by 50%^4^. When *m, P_c_*, and parasite load are known, the parasite growth inhibition (PGI) by the host response at that point in time can be calculated. We determined the relationships between ascending parasite load, onset of fever, *m*, and *P_c_* and their inter-individual variation in 97 malariatherapy subjects in the reference dataset. We then sought the best fit for the distribution of these parameters and their inter-relationships to explain parasite load and duration of illness at the time of presentation using data from 139 Gambian children with malaria (Extended Data Table 1). To provide an additional point of reference we assumed that plasma TNF concentration in the Gambian children should be related to the intensity of the protective host response because this cytokine has been shown to augment parasite growth inhibition by human cells^5,6^. The optimal relationship between TNF and growth inhibition was determined using a maximum-likelihood approach (see Methods and Supplementary Figure 1). Other model assumptions and definitions are shown in Extended Data Table 2. To accommodate biological variation between Gambian children and adult malariatherapy subjects we allowed parameters to be rescaled to improve the fit between the two datasets, resulting in *P_c_* values in the Gambian children being higher than those in the malariatherapy subjects (see Methods). This implies that Gambian children required a higher parasite load than the adult malariatherapy subjects to stimulate a similar host response and is consistent with epidemiological data showing higher fever thresholds in *P. falciparum* infected children than in adults^7^.

After defining the relationships between parasite load, duration of fever, TNF concentration, *m*, and *P_c_* for the Gambian dataset as a whole (Figure 1c), we predicted values of *m* and *P_c_* for individual Gambian children based on their measured parasite load and plasma TNF concentration and their reported duration of fever (Extended Data Table 3). Parasite load, TNF, predicted *m*, and predicted *P_c_* values were highest in those with the most severe manifestations of malaria (SM2), intermediate in those with prostration as the only manifestation of severe disease (SM1), and lowest in uncomplicated malaria (UM) (Fig 1d-g). Duration of illness and estimated PGI at the time of presentation did not differ significantly by clinical phenotype (Fig 1h-i).

Since age can be a major determinant of malaria severity and naturally acquired immunity^8,9^, we examined whether age was associated with *m* or *P_c_*. In this population age was not significantly correlated with *m* but was significantly negatively correlated with *P_c_* (Fig 1j-k). This implies little age-related acquisition of constitutive resistance (for example, naturally-acquired antibody-mediated immunity) in children in this region of The Gambia, which might be expected from the relatively low malaria transmission^10,11^. However, these data also indicate that a lower parasite load should be needed to generate an equivalent protective host response to parasites in older individuals.

To determine whether our model-derived estimates could be used to aid discovery of mechanistic correlates of resistance to *P. falciparum* we performed RNA sequencing on whole blood collected from 23 of the subjects (13 with UM, 10 with SM, Extended Data Table 4) at time of presentation. To avoid confounding by differences in the proportions of major leukocyte subpopulations we performed gene signature-based deconvolution and adjusted total gene expression for cell-mixture. 51 human genes were significantly correlated (26 positively, 25 negatively) with the estimated PGI after adjustment for false discovery rate. We reasoned that these genes should be enriched for effector mechanisms which control parasite load *in vivo*. The positively correlated genes (Fig 2a), associated with enhanced control of parasite growth, showed limited canonical pathway enrichments (Extended Data Table 5) but 16 (62%) were linked together in a network around extracellular signal-regulated kinases ERK1/2 and AKT serine/threonine kinase (Fig 2b). These kinases integrate cellular inflammatory and metabolic responses to control innate defence mechanisms such as cytokine secretion, phagocytosis and degranulation^12,13^. The 25 genes negatively correlated with PGI (Fig 2c), associated with poorer control of parasite growth, were strongly enriched in immune response pathways (Extended Data Table 5). Network analysis showed 15 (60%) of the negatively correlated genes were linked through a network focussed around type 1 interferon (Fig 2d). These findings are consistent with observations that sustained type 1 interferon signalling can impair control of parasite load in mice^14–18^ and potentially in humans^14,19^. C-X-C motif chemokine ligand 10 (CXCL10, also known as IFN-γ-inducible protein of 10 kDa, IP-10) expression had the greatest log-fold change of the genes negatively correlated with PGI (Fig 2c), consistent with findings that CXCL10 expression impairs control of parasite load in mice^20^.

**Fig 2.**
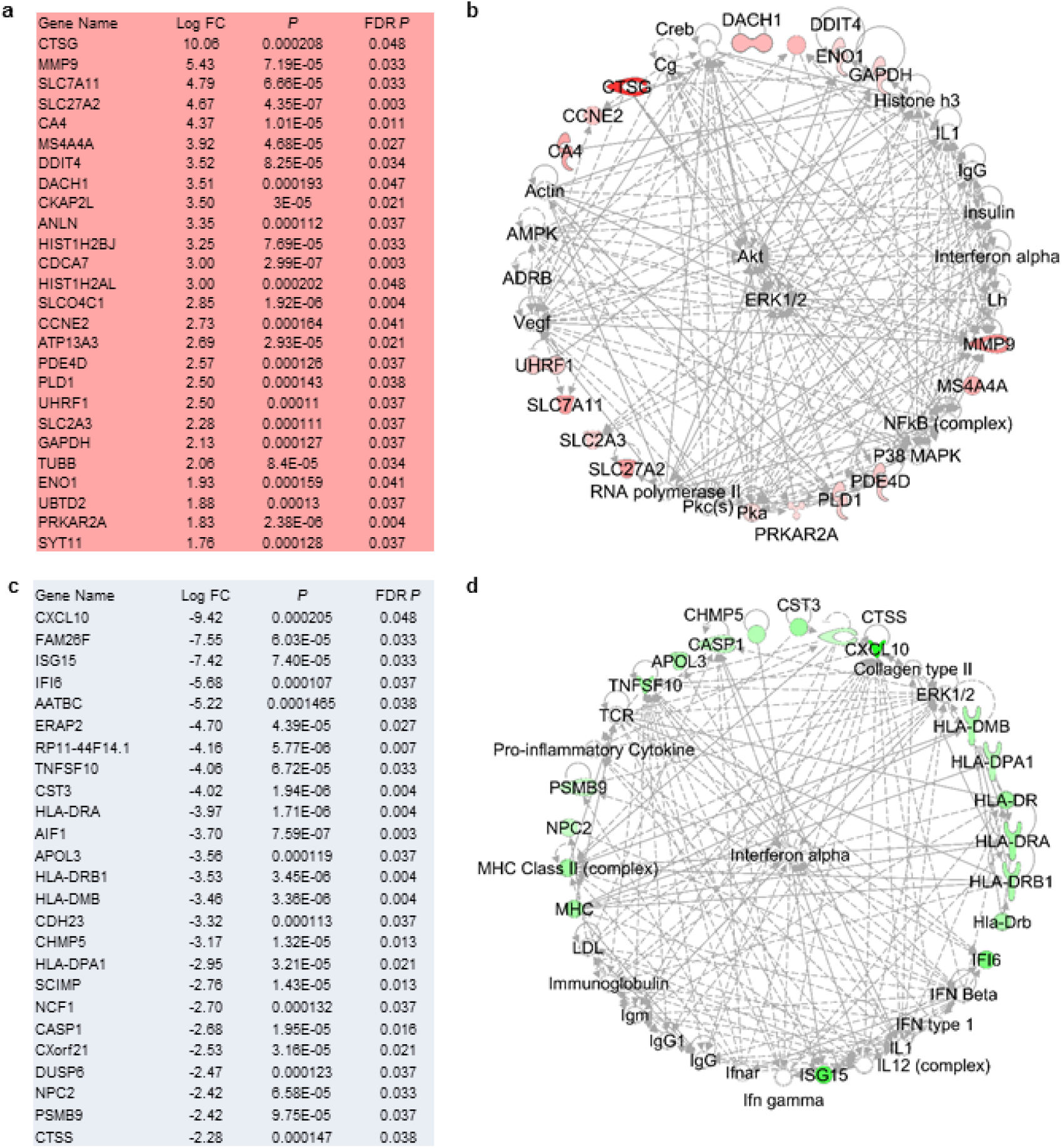
Transcriptional correlates of parasite growth inhibition. (**a, b**) Genes significantly positively correlated with parasite growth inhibition (ie. predicted to reduce parasite growth) after adjustment for false discovery (**a**) and primary network derived from these genes (**b**). (**c, d**) Genes significantly negatively correlated with parasite growth inhibition (predicted to increase parasite growth) after adjustment for false discovery (**c**), and primary network derived from these genes (**d**).

None of the genes positively correlated with PGI have previously been described as mediators of resistance to malaria so we sought direct biological evidence, focussing on genes encoding secreted proteins as the best candidates. Of the 26 genes, two encode predominantly secreted proteins: *CTSG* (cathepsin G) and *MMP9* (matrix metallopeptidase 9, also known as gelatinase B). Cathepsin G localizes in neutrophil azurophil granules whilst MMP9 localises in neutrophil gelatinase and specific granules^21^. Treatment with recombinant cathepsin G and MMP9 inhibited growth of *P. falciparum* 3D7 strain *in vitro* (Fig 3a). In order to elucidate mechanisms of action we examined whether these proteases inhibited parasite growth by preventing invasion of erythrocytes. Addition of cathepsin G to schizont cultures produced a dramatic reduction in invasion, and pretreatment of erythrocytes with cathepsin G before adding them to schizont cultures produced a similar reduction in their invasion (Fig 3b), indicating that cathepsin G acts primarily on the erythrocyte (Fig 3b). Addition of MMP9 to schizont cultures produced a more modest reduction, whilst pretreatment of erythrocytes did not reduce invasion, implying that MMP9 likely acts against schizonts or free merozoites rather than preventing invasion at the erythrocyte surface (Fig 3b).

In order to identify biologically relevant concentrations of cathepsin G and MMP9 we measured their concentrations in whole blood from healthy donors, before and after degranulation was stimulated with PMA, and in plasma from children with malaria at the time of clinical presentation (Fig 3c). Local concentrations which might occur *in vivo*, adjacent to degranulating neutrophils, could be at least an order of magnitude higher^22^. MMP9 dose-dependently inhibited parasite growth over a physiological range of concentrations (Fig 3d). Similarly, parasite invasion was dose-dependently inhibited by cathepsin G pre-treatment of erythrocytes, with similar effects in each of four parasite strains with different invasion phenotypes^23^ (Fig 3e). We therefore asked whether cathepsin G might cleave a range of RBC surface proteins which are used as invasion receptors by *P. falciparum* ^24^. Consistent with its broad inhibition of parasite invasion, cathepsin G dose-dependently cleaved the majority of *P. falciparum* invasion receptors including glycophorins A, B, and C, CD147 (basigin), CD108 (semaphorin 7A), and complement receptor 1 (CR1), but not CD55 (DAF) (Fig 3f). MMP9 did not cleave any of these surface receptors (Extended Data Figure 2). PMA stimulation of healthy donor whole blood recapitulated the loss of erythrocyte surface glycophorins A and B, CD108 and CD147 in all donors, decreased glycophorin C expression in 6 of 8 healthy donors, but did not consistently reduce CR1 (Fig 3g) (as might be expected from the dose-response curves, Fig 3f). In samples from Gambian children on the day of presentation with *P. falciparum* malaria, the proportions of erythrocytes with detectable expression of glycophorins A and B and CD147 were significantly lower than in convalescent samples (28 days after treatment), and there was a trend to lower expression of CD108 and glycophorin C (Fig 3h). These results would be consistent with cleavage of these surface molecules *in vivo* during acute infection. The variable expression seen at day 28 (Fig 3h) may indicate the persistence of modified erythrocytes in the circulation. The importance of glycophorins and basigin in RBC invasion and genetic susceptibility to severe malaria is well established^25–28^, and so it is highly likely that the cleavage of these erythrocyte receptors by cathepsin G would have a protective effect *in vivo*.

**Fig 3.**
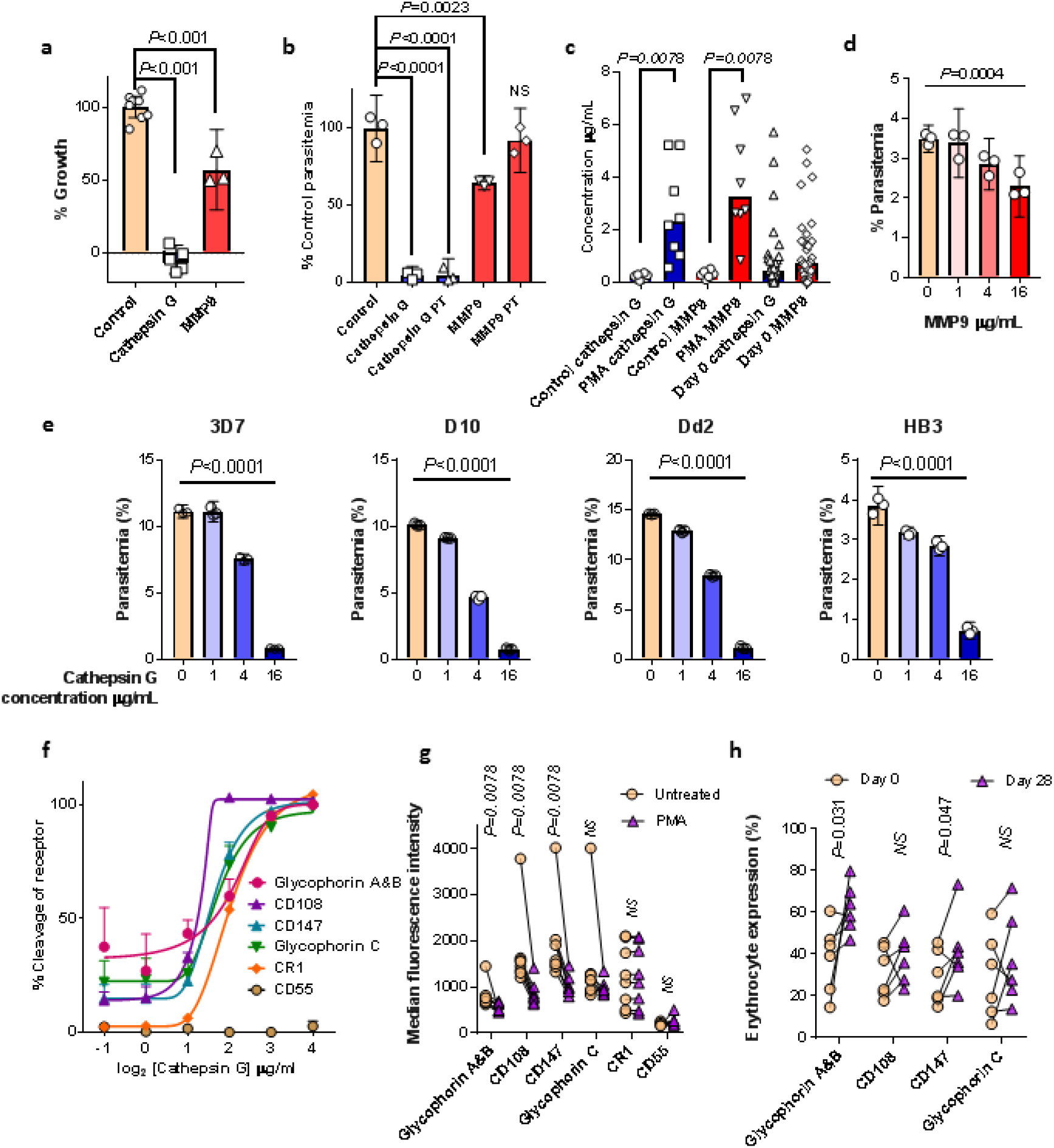
Effects of cathepsin G and MMP9 on parasite growth and expression of erythrocyte invasion receptors. (**a**) Effect of cathepsin G (18µg/mL) and MMP9 (16µg/mL) on *in vitro* growth (n=3–8 biological replicates, representative of two independent experiments). (**b**) Effect of cathepsin G (18µg/mL) and MMP9 (18µg/mL) on erythrocyte invasion of *P. falciparum* 3D7 when added directly to schizonts and donor red cells, or when pre-incubated (PT) with donor red cells before washing and adding to schizonts (n=3 biological replicates per condition, representative of two independent experiments). (**a**,**b**) Show mean (95% CI) and *P* for unpaired t-test. (**c**) Cathepsin G and MMP9 concentrations in plasma from healthy donor whole blood (n=8) unstimulated or stimulated with 1µM PMA, and from Gambian children with *P. falciparum* malaria (n=34). Bars show median, *P* for Wilcoxon matched pairs test. (**d, e**) Dose response for growth inhibition by MMP9 against *P. falciparum* 3D7 and (**d**) and for invasion inhibition using cathepsin G pre-treated RBCs against four parasite strains (n=3 biological replicates per dose, mean (95% CI) and *P* for linear trend, representative of two independent experiments). (**f**) Dose response for erythrocyte surface receptor cleavage by cathepsin G (n=3 biological replicates per dose, mean +/- standard error, asymmetric 5-parameter logistic regression fit lines, representative of two experiments). (**g**) Effect of PMA stimulation of healthy donor (n=8) whole blood on erythrocyte surface receptor expression assessed by fluorescence intensity (*P* for Wilcoxon matched pairs test). (**h**) Comparison of proportion of erythrocytes with detectable receptor expression in acute (day 0) and convalescent (day 28) samples from Gambian children with malaria (n=6, *P* for one-sided Wilcoxon test).

Having shown that mechanisms of host resistance can be identified from their correlation with PGI, we asked whether we could also identify mechanisms controlling *m*. In our model, *m* may be influenced by constitutive factors but should be independent of any parasite load-dependent host response. Therefore we sought to confirm expected associations with two constitutive host factors known to influence parasite growth: iron^29^ and complement factor H (FH)^30,31^. Since we did not have premorbid blood samples we used convalescent blood, collected 28 days after treatment, as a proxy for pre-infection status (Supplementary Dataset). Iron deficiency is protective against malaria^32^ and reduces parasite multiplication *in vitro* ^29^. Consistent with this, levels of hepcidin (a regulator of iron metabolism and marker of iron sufficiency or deficiency^33^) were significantly correlated with *m* (r=0.27, *P*= 0.009) in 92 children who had not received blood transfusion (Extended Data Figure 3). FH is a constitutive negative regulator of complement activation which protects host cells from complement mediated lysis^34^. Many pathogens including *P. falciparum* have evolved FH binding proteins^30,31^, and FH protects blood-stage parasites from complement mediated killing *in vitro* ^30,31,34^. In 14 children with additional day 28 plasma available (Supplementary Data Set) we found that FH was significantly correlated with *m* (r=0.66, *P* = 0.01) (Extended Data Figure 3). Thus, the quantitative estimates of parasite multiplication and PGI from our model exhibit the expected relationships with known constitutive determinants of parasite growth and this provides further evidence to support the validity of the method.

Using a model-based approach to estimate the within-host dynamics of pathogen load and its determinants in human infection we could estimate the relative contributions of parasite multiplication and host response to pathogen load measured at a single point in time, and we have used these predictions to identify mechanistic correlates of host resistance to malaria. Our approach is based on assumptions which are reasonable, but largely unverifiable, and alternative approaches are possible. However, our biological validation suggests that the relative estimates of *m* and PGI are accurate enough to be useful, providing proof-of –principle that pathogen load dynamics can be estimated in humans. Our approach could be refined and expanded to identify additional biological determinants of pathogen load such as genetic correlates of *m* and *P_c_*, or serological correlates of *m* and PGI. This has the potential to yield further novel insights into host-pathogen interactions in malaria, to facilitate discovery of new therapeutic and vaccine strategies, and improve predictive modelling of their impact on disease. Our findings indicate leads for development of host-directed therapeutics to augment antimalarial treatment, particularly in the setting of drug resistant parasites. Inhibition of type-1 interferon or CXCL10 signalling is a realistic option with inhibitory antibodies and small molecules already in development for other indications^35,36^. The therapeutic potentials of cathepsin G and MMP9 may be counterbalanced by risk of collateral tissue damage, but selective targeting of receptors on the erythrocyte surface may be a useful paradigm for both treatment and prevention of malaria.

**Extended Data Table 1.**
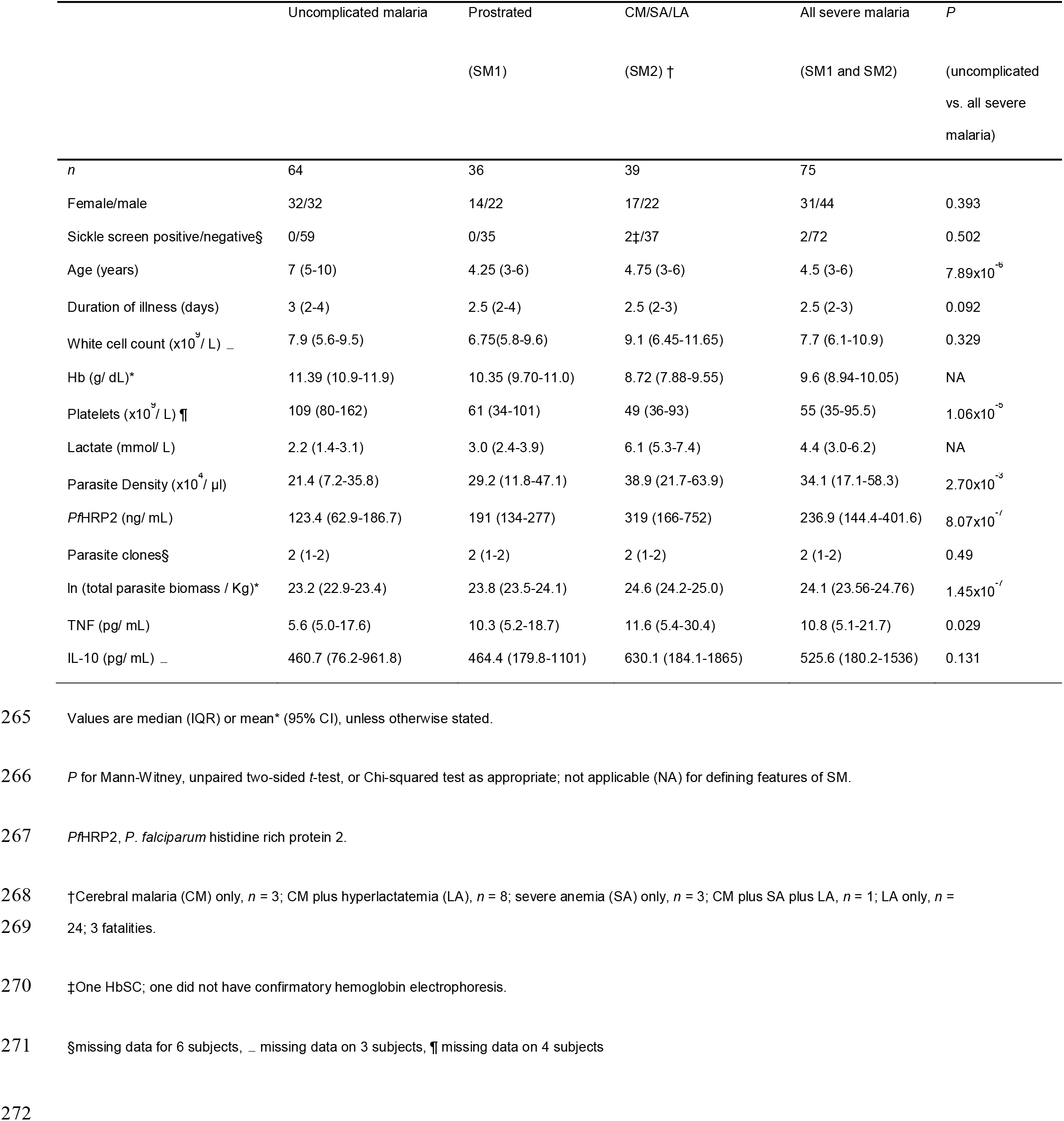
Characteristics of 139 Gambian children with malaria.

**Extended Data Table 2.**
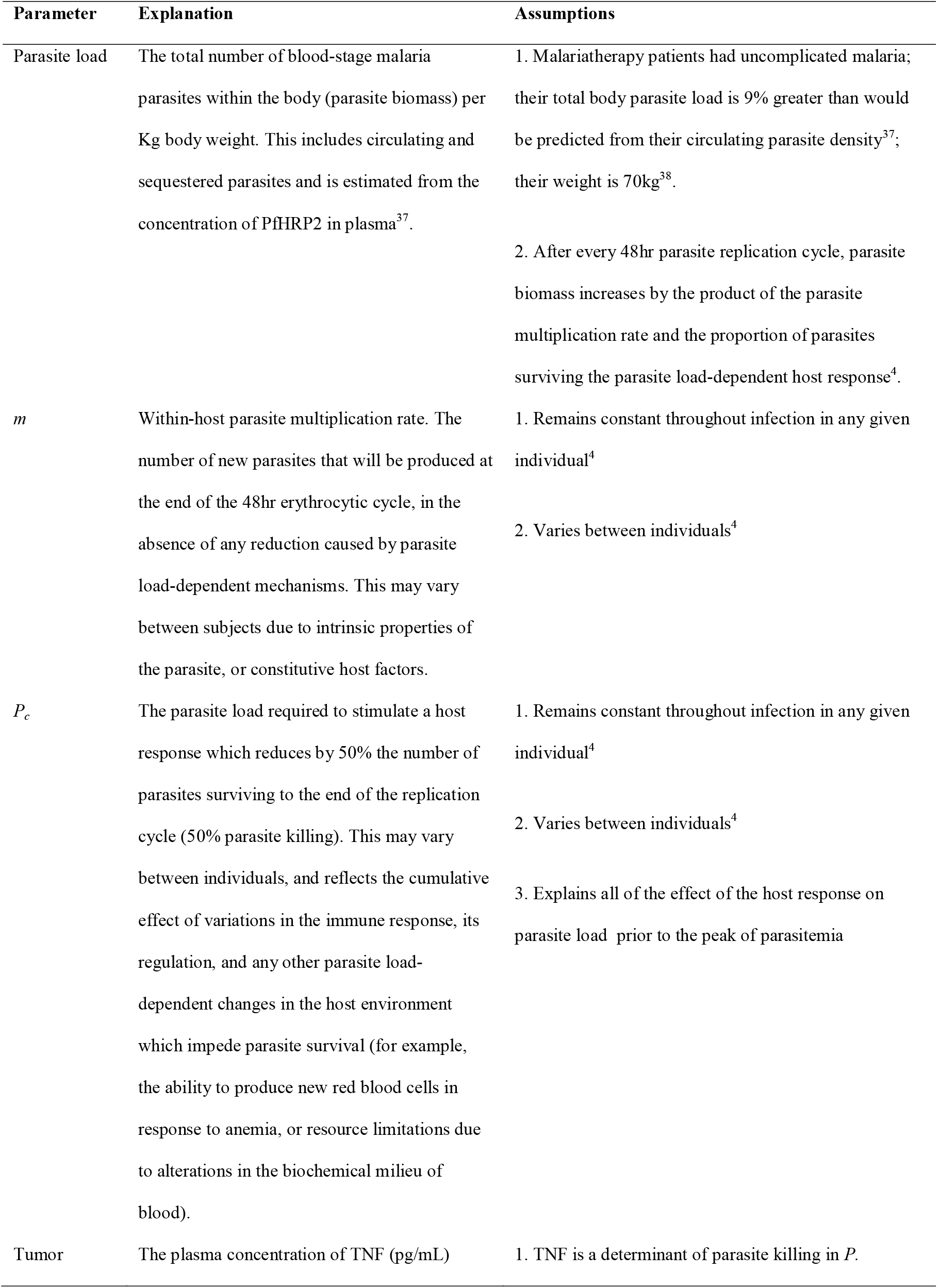
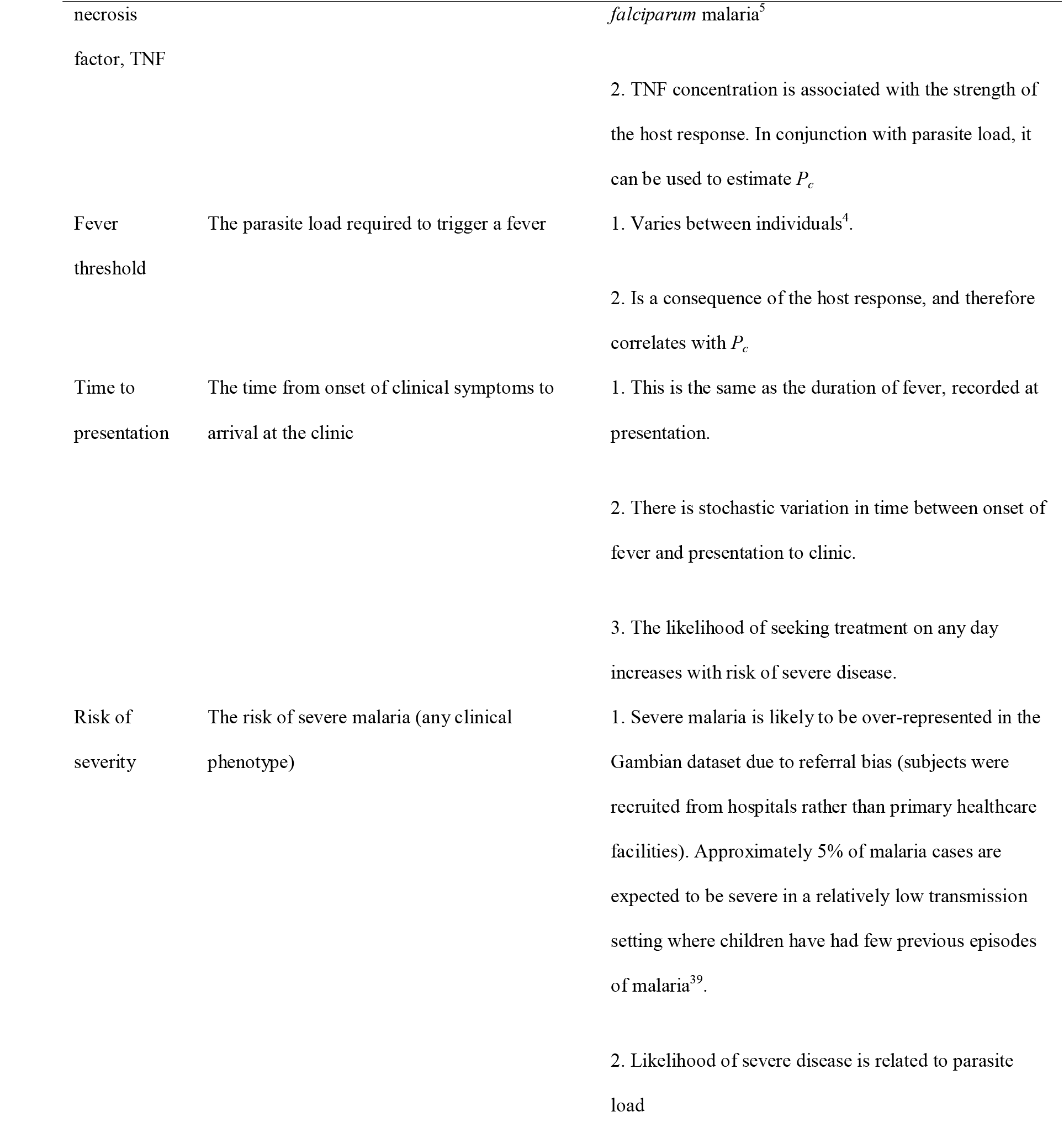
Model parameters and assumptions.

**Extended Data Table 3.**
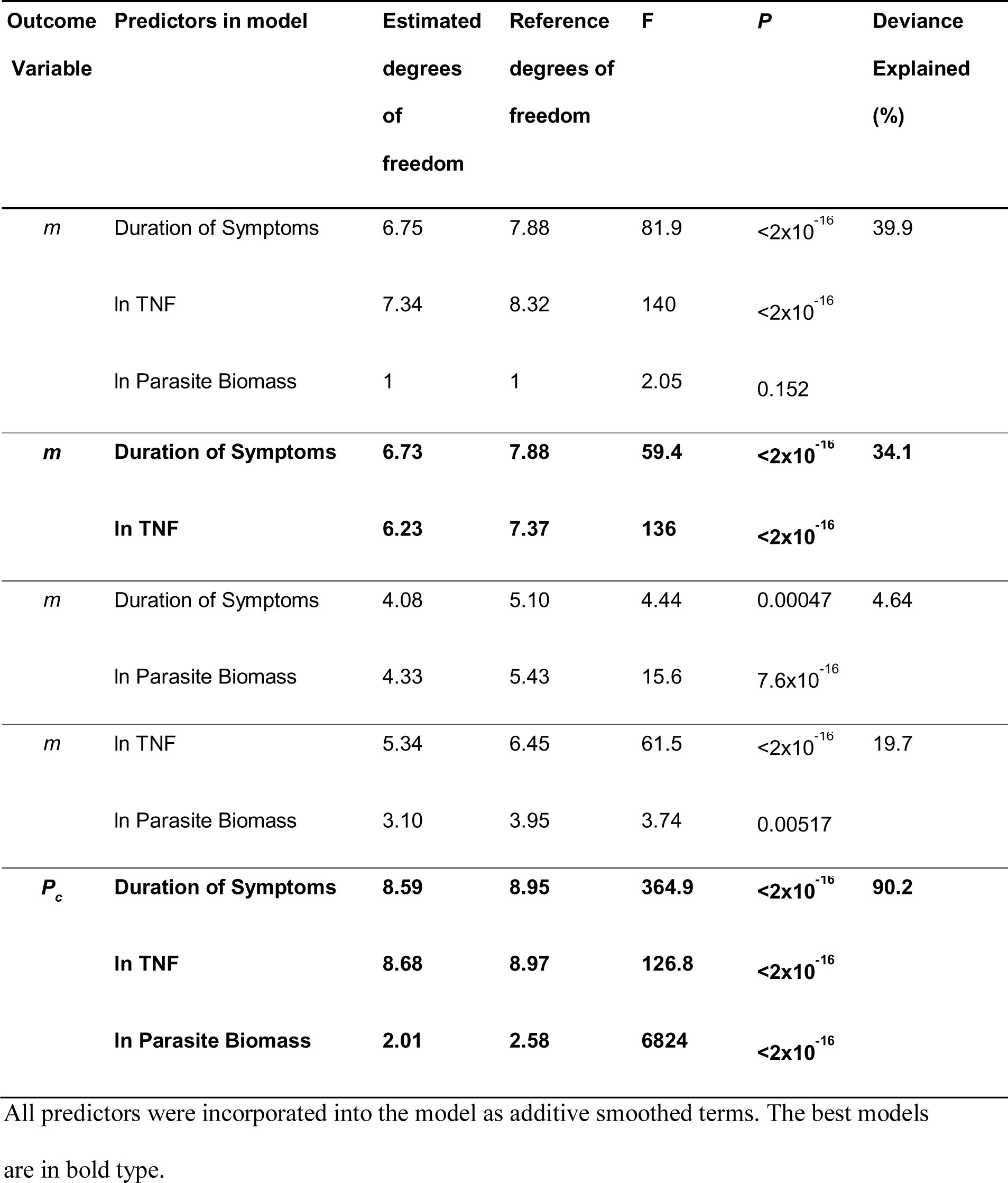
Comparison of general additive models to predict *m* and *P_c_*.

| Outcome Variable | Predictors in model | Estimated degrees of freedom | Reference degrees of freedom | F | P | Deviance Explained (%) |
| --- | --- | --- | --- | --- | --- | --- |
| *m* | Duration of Symptoms | 6.75 | 7.88 | 81.9 | $<2 \times 10^{-16}$ | 39.9 |
| | In TNF | 7.34 | 8.32 | 140 | $<2 \times 10^{-16}$ | |
|  | In Parasite Biomass | 1 | 1 | 2.05 | 0.152 |  |
| ***m*** | **Duration of Symptoms** | **6.73** | **7.88** | **59.4** | **$<2 \times 10^{-16}$** | **34.1** |
|  | **In TNF** | **6.23** | **7.37** | **136** | **$<2 \times 10^{-16}$** |  |
| *m* | Duration of Symptoms | 4.08 | 5.10 | 4.44 | 0.00047 | 4.64 |
| | In Parasite Biomass | 4.33 | 5.43 | 15.6 | $7.6 \times 10^{-16}$ | |
| *m* | In TNF | 5.34 | 6.45 | 61.5 | $<2 \times 10^{-16}$ | 19.7 |
|  | In Parasite Biomass | 3.10 | 3.95 | 3.74 | 0.00517 |  |
| ***P<sub>c</sub>*** | **Duration of Symptoms** | **8.59** | **8.95** | **364.9** | **$<2 \times 10^{-16}$** | **90.2** |
|  | **In TNF** | **8.68** | **8.97** | **126.8** | **$<2 \times 10^{-16}$** |  |
|  | **In Parasite Biomass** | **2.01** | **2.58** | **6824** | **$<2 \times 10^{-16}$** |  |

All predictors were incorporated into the model as additive smoothed terms. The best models are in bold type.

**Extended Data Table 4.** Characteristics of subjects used for RNA sequencing.

|  | UM | SM |
| --- | --- | --- |
| *n* | 13 | 10 |
| Female/male | 5/8 | 3/7 |
| Sickle screen positive/negative | 0/12 | 1<sup>§</sup>/9 |
| Age (years) | 8 (4-10) | 4.6 (3.3-5) |
| Duration of illness (days) | 2 (2-3) | 2.25 (2.0-2.9) |
| White cell count (x10<sup>9</sup> / L) | 8.6 (7.3-10.6) | 10.2 (8.8-11.6) |
| Hemoglobin (g/ dL) | 10.8 (10.5-12.4) | 7.8 (6.8-9.8) |
| Platelets (x10<sup>9</sup> / L) | 123 (104-132) | 53 (41-114) |
| Lactate (mmol/ L) | 2.2 (1.6-3.0) | 5.8 (4.4-10.3) |
| Parasite Density (x10<sup>4</sup> / µl) | 23 (20.8-35.2) | 24.1 (13.6-27.9) |
| *Pf*HRP2 (ng/ mL) | 163 (128-187) | 589 (209-2030) |
| Parasite clones* | 2 (1-2) | 1 (1-2) |
| ln (total parasite biomass / Kg) | 23.4 (23.3-23.7) | 25.1 (24.1-26.3) |
| TNF (pg/ mL) | 5.0 (5.0-12.1) | 11.2 (5.6-16.0) |
| IL-10 (pg/ mL) | 651 (252-1900) | 429 (201-1090) |
| Estimated ln *P<sub>c</sub>* | 11.8 (10.9-12.0) | 13.3 (11.9-13.5) |
| Estimated *m* | 12.5 (12.5-14.2) | 13.5 (12.3-15.6) |
| Estimated PGI | 0.69 (0.55-0.75) | 0.71 (0.57-0.82) |

For continuous data values are median (interquartile range). <sup>§</sup>HbSC, \*data available for 11 UM and 9 SM.

**Extended Data Table 5.** Most enriched gene ontology (GO) biological process terms

| GO term | Description | Number of<br>genes in<br>background | Number of<br>genes<br>positively<br>correlated<br>with PGI | Enrich<br>ment *P* | Number of<br>genes<br>negatively<br>correlated<br>with PGI | Enrich<br>ment *P* |
| --- | --- | --- | --- | --- | --- | --- |
| GO:0015711 | organic anion<br>transport | 183 | 4 | **0.0001**<br>**7** | 1 | 0.47 |
| GO:0061615 | glycolytic<br>process through<br>fructose-6-phosphate | 18 | 2 | **0.0003**<br>**6** | 0 | 1 |
| GO:0035606 | peptidyl-<br>cysteine S-trans-<br>nitrosylation | 2 | 1 | **0.0032** | 0 | 1 |
| GO:0035092 | sperm<br>chromatin<br>condensation | 2 | 1 | **0.0032** | 0 | 1 |
| GO:0051186 | cofactor<br>metabolic<br>process | 243 | 3 | **0.0064** | 0 | 1 |
| GO:0002396 | MHC protein<br>complex<br>assembly | 5 | 0 | 1 | 4 | **5.7 x10<sup>-</sup>**<br>**10** |
| GO:0034097 | response to<br>cytokine | 523 | 1 | 0.58 | 11 | **7.17**<br>**x10<sup>-7</sup>** |
| GO:0045087 | innate immune<br>response | 747 | 4 | 0.028 | 12 | **3.41**<br>**x10<sup>-6</sup>** |
| GO:0002376 | immune system<br>process | 1733 | 4 | 0.30 | 17 | **1.33**<br>**x10<sup>-5</sup>** |
| GO:0032649 | regulation of<br>interferon-<br>gamma<br>production | 54 | 0 | 1 | 3 | **0.00080** |

The five most enriched, non-redundant biological process terms were determined using
REVIGO for genes positively or negatively associated with parasite growth inhibition

**Extended Data Table 6.** Antibodies used to assess erythrocyte surface receptor expression

| Antibody, Clone | Isotype control |
| --- | --- |
| APC Mouse anti-human CD235ab Antibody, Clone HIR2 (Biolegend) | APC Mouse IgG2b, κ (Biolegend) |
| PE Mouse anti-human CD108 Antibody, Clone MEM-150 (Biolegend) | PE Mouse IgM, κ (Biolegend) |
| Alexa Fluor® 488 Mouse anti-human CD147 Antibody, Clone HIM6 (Biolegend) | Alexa Fluor® 488 Mouse IgG1, κ (Biolegend) |
| Alexa Fluor® 405 Mouse anti-human Glycophorin C Antibody (BRIC10) (Novus Biologicals) | Alexa Fluor® 405 Mouse IgG1 (Novus Biologicals) |
| PE Mouse anti-human CD35 Antibody (CR1), Clone E11 (Biolegend) | PE Mouse IgG1 (Biolegend) |
| FITC Mouse anti-Human CD55 Clone IA10 (RUO) (BD Biosciences) | FITC Mouse IgG2a, κ (BD Biosciences) |

## Extended Data Figures

**Extended Data Figure 1.**
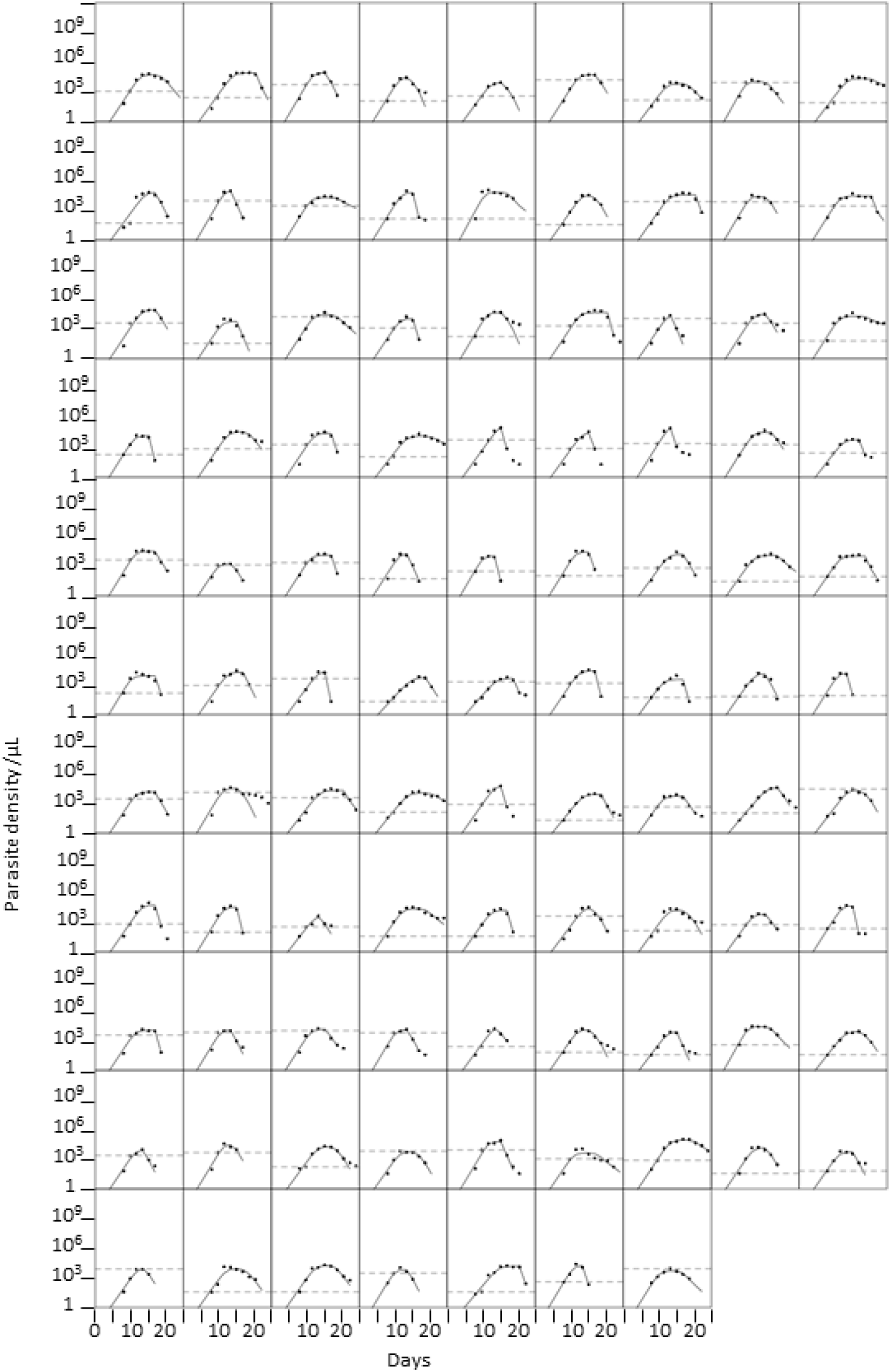
The dynamics of pathogen load in malariatherapy patients. Change in parasite density over time (for the first wave of parasitemia in 97 malariatherapy patients. Each plot represents one subject. Dots indicate observed parasite densities on alternate days; dashed line, observed fever threshold: solid line: fit of modelled trajectory of parasite density.

**Extended Data Fig. 2.**
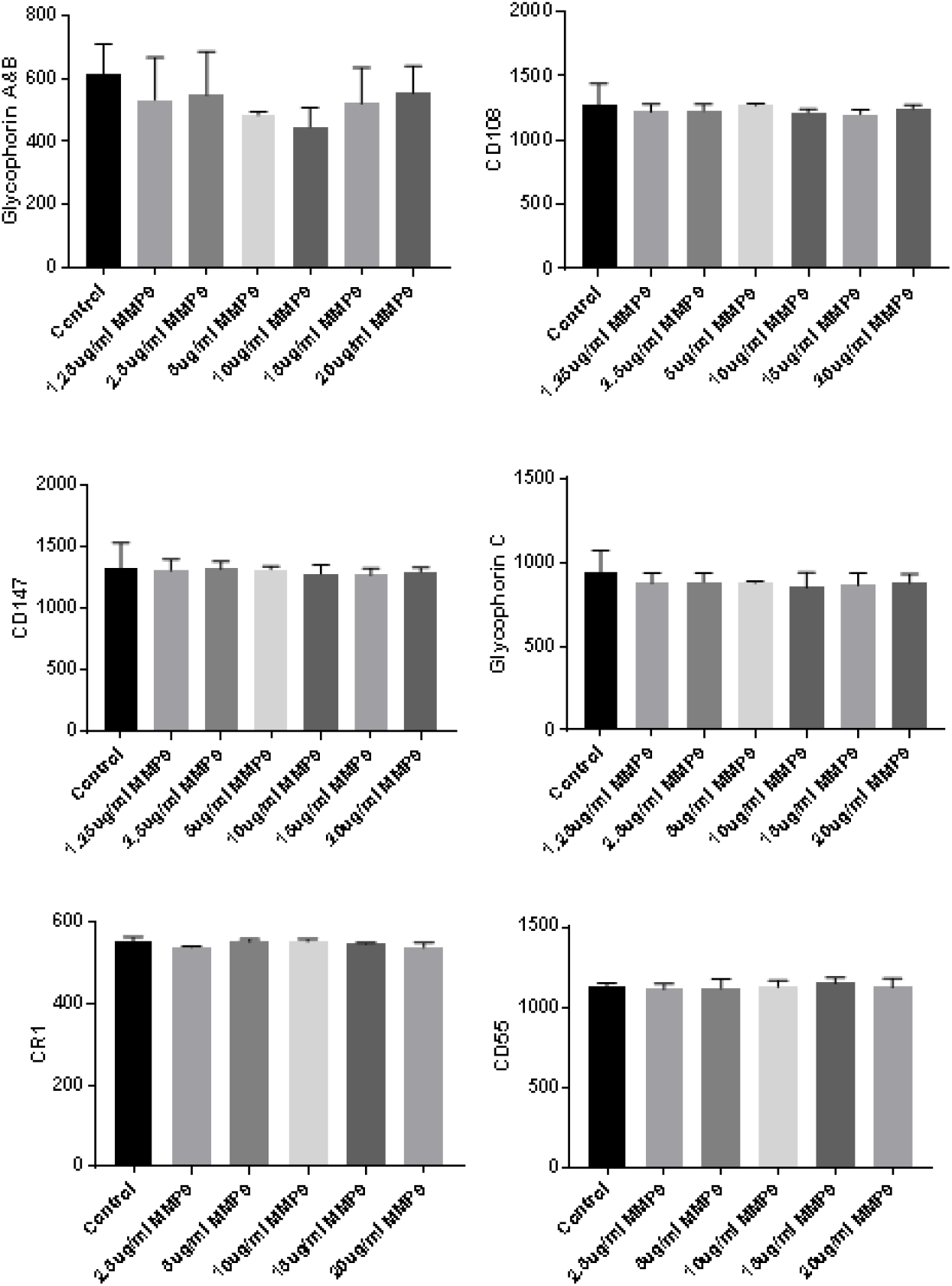
Effect of MMP9 treatmenton erythrocyte surface receptors. Median fluorescent intensity of erythrocyte surface molecule expression following MMP9 treatment (n=3 biological replicates per condition, mean and range). None were significantly different to the control condition using t-test.

**Extended Data Fig 3.**
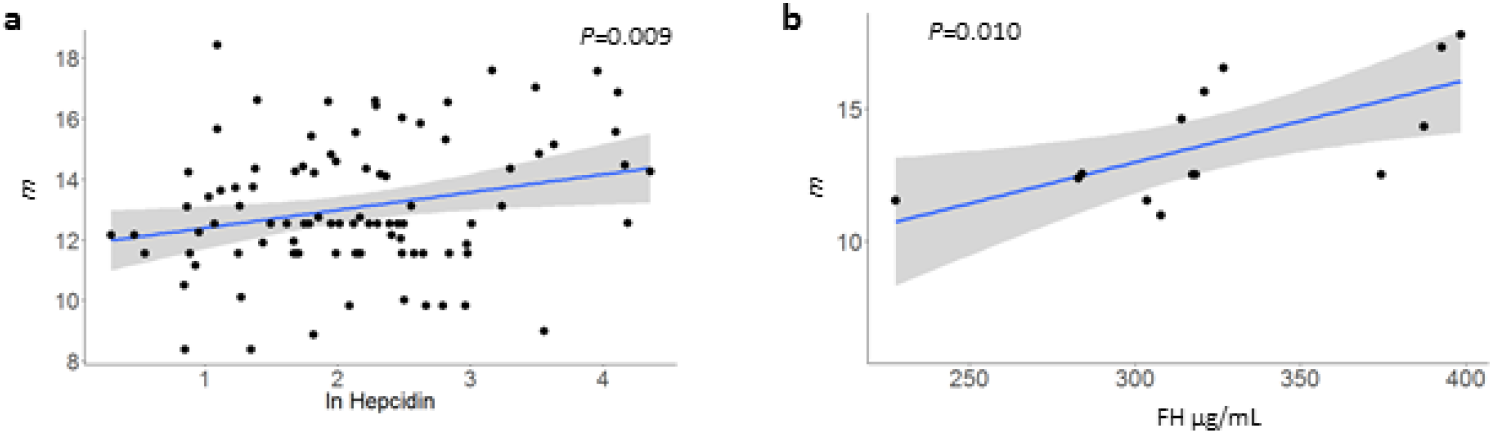
Correlates of parasite multiplication rate. (**a, b**) Predicted parasite multiplication rate, *m*, was correlated with hepcidin concentration (**a**, n=92) and complement factor H (FH) concentration (**b** n=14) in convalescent plasma obtained from children 28 days after treatment for malaria. Blue line, linear regression: grey shading. 95% Cl; *P*, Pearson correlation.

**Extended Data Fig. 4.**
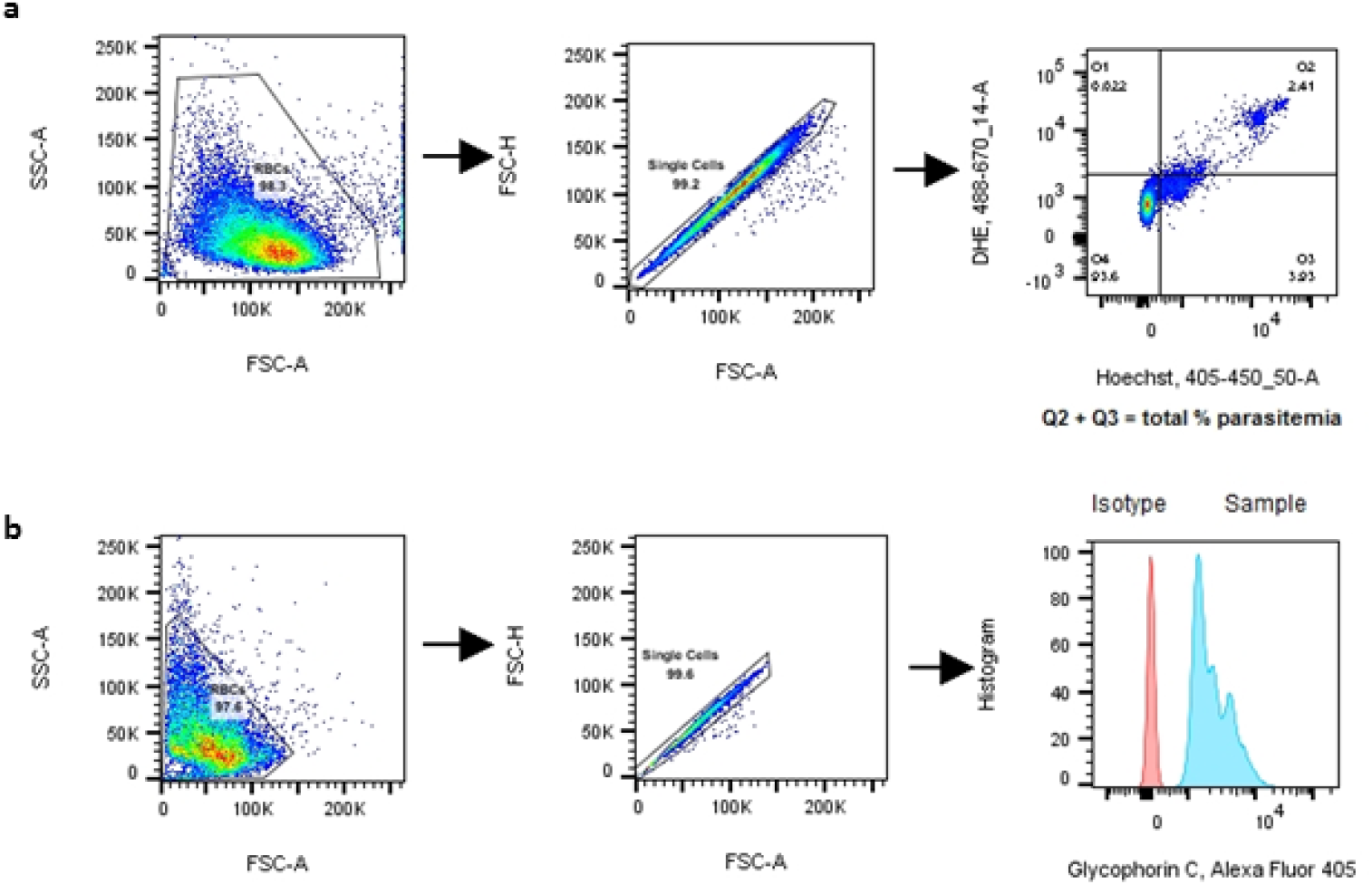
Gating strategies fortlow cytometry analyses. (**a**) Gating for determination of parasitemia by flow cytometry based on staining of infected red blood cell with dihydroethidium (DHE) and Hoechst. (*b*) Gating on red blood cells for determination of surface molecule expression by fluorescence intensity (example of glycophorin C fluorescence compared with isotype fluorescence).

## Materials and Methods

### Subjects and laboratory assays

We used data from all of the malariatherapy patients reported by Dietz et al.^4^ and from all 139 Gambian subjects reported in our previous studies^37,40,41^ who had all of the following data available: age, parasite biomass estimate, plasma TNF concentration, duration of illness and severity of illness. No subjects were excluded after this selection, and all available data was included in analyses, without any exclusion of outliers. As described previously^37,40,41^, Gambian children (<16 years old) were recruited with parental consent from three peri-urban health centres in the Greater Banjul region, from August 2007 through January 2011 as part of a study approved by the Gambia Government/MRC Laboratories Joint Ethics Committee, and the Ethics Committee of the London School of Hygiene and Tropical Medicine. *P. falciparum* malaria was defined by compatible clinical symptoms in the presence of ≥5000 asexual parasites/µL blood, and any children suspected or proven to have bacterial co-infection were excluded. Severe malaria was specifically defined by the presence of prostration (SM1) or any combination of three potentially overlapping syndromes (cerebral malaria (CM), severe anemia (SA, hemoglobin <5 g/dL), and hyperlactatemia (blood lactate >5 mmol/L) – collectively SM2)^37,41,42^. Clinical laboratory assays, measurements of plasma TNF and IL-10 by Luminex, measurements of gene expression by RT-PCR, and estimation of total parasite biomass from *Pf*HRP2 ELISA have been previously described^37,41^. Subject-level data from these Gambian children is available as **Supplementary Dataset**.

### Statistical analyses

Statistical analyses were undertaken using the R statistical software^43^ and GraphPad Prism (GraphPad Software, Inc.). Continuous variables were compared between groups using unpaired or paired student’s t-test (when normally distributed) and the Mann-Whitney-Wilcoxon or Wilcoxon matched pairs tests (when normal distribution could not be assumed), or ANOVA for comparison across multiple groups. Correlations were assessed using Pearson’s correlation, after log transformation of the data if appropriate. All hypothesis tests were two-sided with alpha = 0.05, unless specifically indicated otherwise. One-sided testing was only used when consistent with the *a priori* hypothesis. Associations between explanatory variables and latent variables were assessed using generalized additive models (GAM^44^, using the R package “mgcv”) and explained variance was used to select the best GAM once all model terms were significant (*P*<0.05). Dose-response curves were fitted using asymmetrical sigmoidal five-parameter logistic equation in GraphPad Prism.

### Model relating parasite multiplication, host response and parasite load

A process-based, stochastic simulation model was devised to reproduce the clinical data collected from the Gambian children. This was achieved by combining the information in the Gambian data with a model describing the first wave of parasitemia in non-immune adults who were deliberately infected with *P. falciparum* malaria to treat neurosyphilis (“malariatherapy”)^4^. These malariatherapy data, from the pre-antibiotic era, are the main source of information on the within-host dynamics and between-host variation in the course of parasitemia in untreated malaria infections. The model of Dietz et al.^4^ was modified and extended in order to be applied to the Gambian data.

*Model of ascending parasitemia in malariatherapy subjects*. The model relates parasite density after each 2-day asexual blood stage cycle (P_(t+2)_) to the parasite density at the end of the previous cycle (P_(t)_) by the following equation:

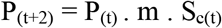

The host-specific parasite multiplication rate, *m*, is a property of both parasite and host, allowing for growth-inhibition by constitutive factors; the proportion of parasites that will survive the effects of the density-dependent host response in the present cycle is S_c_:

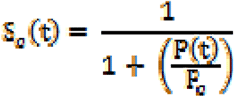

where *P_c_* is the host-specific parasite load threshold at which the host response is strong enough to inhibit 50% of parasite growth in that cycle. Consistent with the original Dietz model, P_(0)_ was set to 0.003 parasites/µl^4^.

The original Dietz model included an additional parameter, S_*m*_, to help describe the decline in parasitemia after the peak of the first wave. S_*m*_ is the proportion of isogenic parasites surviving an additional density- and time-dependent host response, which might represent adaptive immunity^4^. Estimates of the range of values of S_*m*_ in the Dietz dataset and model were used when simulating data but since this parameter has little influence on parasite densities prior to the peak it was not used to make subsequent predictions of *m* and *P_c_* in individual Gambian subjects.

At the explicit request of Klaus Dietz and Louis Molineaux, we hereby communicate the following correction regarding their assertion that the malariatherapy patients had not received any treatment^4^: it was later found that 47 of these patients had indeed received subcurative treatment, and that those patients had significantly higher parasite densities. This is unlikely to influence our analysis, because treatment would only be provided when malariatherapy patients became very unwell, presumably at maximum parasitemia, whereas most patients with naturally acquired infection likely present prior to the peak parasitemia that might occur in the absence of treatment.

*Fitting of the malariatherapy model to data from Gambian children.* Individual-level parameter estimates for the malaria therapy dataset were kindly provided by Klaus Dietz. The logarithms of these 97 estimates of *m* and *P*_c_ were well described by a multivariate normal distribution, providing a quantitative description of inter-individual variation in the dynamics of the first wave of parasitemia. In order to use the Dietz model to simulate the Gambian data, a number of modifications and extensions were made. Some of these required estimation of additional parameters by comparing the model simulations with the Gambian data. Dietz et al. provided a statistical description of the parasite density at which first fever occurred (the “fever threshold”) in the form of the distribution of the ratio of threshold density to peak parasitemia. The median density at first fever was at 1.4% of peak density. We introduced the assumption that the onset of fever occurs at a particular threshold value of *S_c_*, because fever is dependent on the production of cytokines like interleukin-6 and TNF, both components of the host response. This constitutes a process-based model for the onset of fever rather than a purely statistical one. Because individuals differ in their response to parasite load (captured through variation in *P_c_*), this results in variation of parasite densities at first fever but ignores any potential variation among individuals with respect to host response at onset of fever. The host response threshold for the onset of fever 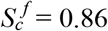 = 0.86 was determined as the value of *S_c_* calculated at 1.4% of the peak density of a simulated individual with the median parameter values. This yielded a distribution of fever ratios similar to the one described by Dietz et al.^4^, albeit with less variation.

To simulate the time between onset of fever and clinical presentation we made use of the self-reported duration of symptoms in the Gambian data. The model which was most consistent with these values assumed a gamma-distributed duration of symptoms in non-severe cases, and a possibility to present earlier in the case of more severe disease. Since parasite biomass is related to likelihood of having severe malaria^37,38^ the probability of early presentation on any day after onset of fever was set proportional to the (density-dependent) probability of having severe disease on that day. Scale (ζ) and shape (κ) parameters of the gamma distribution as well as the factor (ξ) for determining the probability of early presentation were estimated from the Gambian data.

We assumed that TNF production τ(t) increases monotonically with density dependent host response (1-*S_c_*) and represented this relationship using a heuristic function of the form

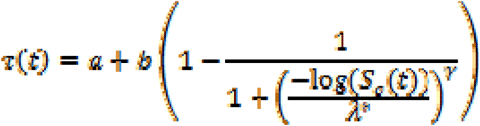

, with free parameters a, b, λ^*^ and γ estimated from the Gambian data.

The Gambian children had on average higher parasite densities than the malariatherapy patients, which led to a bad fit of the original model to the Gambian data. This was overcome by introducing the assumption that the Gambian children had a different range of values of *P_c_* to the adult malariatherapy patients. A factor π was therefore estimated by which the ln *P_c_* value from the Dietz model was multiplied. This led to overall higher parasite densities upon presentation. However, our model uses parasite biomass and its relationship with disease severity to predict day of presentation, and there is an interaction between the mean ln *P_c_* and the variation in ln *P_c_*, as well as the proportion of severe malaria in the simulated Gambian population. Based on the relatively low malaria transmission in the Banjul area of The Gambia, we assumed that severe cases (defined by the presence of any of: prostration, hyperlactatemia, severe anemia or cerebral malaria) were over-represented by hospital-based recruitment and that in an unselected population of children of similar age to those in our dataset only approximately 5% of all malaria infections would be severe^39^. Therefore we estimated a factor δ by which the variance of ln *P_c_* should be multiplied such that both rate of severity as well as the distribution of parasite biomass matched well after fitting our simulation to the Gambian data.

The free parameters ζ, κ, ξ, a, b, λ^*^, γ, π and δ, together summarized as θ, were estimated by fitting model simulations to the information on TNF, parasite density, and duration of symptoms, for any given candidate parameterization, a total of 139 clinically presenting individuals were simulated from the model, which corresponds to the size of the Gambian dataset. An objective function L(θ) was calculated, and a simulated annealing algorithm (provided by the “optim” function in R) determined the value for θ which maximizes this function. The log-likelihood L (θ) was comprised of three separate objectives. The first objective represented the log-probability that the frequency of severe cases in the simulation was equal to an assumed 5%, employing a binomial likelihood, given the actual number of severe cases sampled in 139 simulated individuals. The second objective considered the overlap between the bivariate distribution of ln parasite density vs. ln TNF concentration in the simulated data compared to the Gambian dataset. An approximate numerical value for this partial log-likelihood was obtained as the log probability of the Gambian data (density and TNF) given a two-dimensional kernel density estimate of the simulation output as a likelihood model. Kernel density estimates were obtained using the “kde2d” function in the “MASS” package in R. In this calculation, the TNF/density data points of severe or prostrated Gambian patients entered the partial likelihood with a weight of 1/11, to account for the oversampling of severe cases in the Gambian data. The third objective concerned the two- dimensional distribution of log density and duration since first fever. This partial log-likelihood was obtained using the same kernel-based approach described above, with weights of 1/11 for severe and prostrated cases. The overall log-likelihood L (θ) was calculated as a weighted sum of the three partial log-likelihoods, with the log-probability of having the desired true severity rate weighted with a factor of 68, which ensured similar magnitude of the three partial log-likelihoods at the optimum.

The results of the fitting algorithm were visually confirmed to yield a good overlap of the joint distributions of density and biomass, the duration of symptoms, TNF and biomass between simulation and the Gambian children. Approximate confidence intervals for the parameter estimates were determined by employing a 2^nd^ degree polynomial to estimate the curvature of the maximum simulated likelihood surface in the vicinity of the parameter point estimate, assuming independence of parameters.

As in the original model of Dietz et al.^4^, peripheral parasite densities were used to determine the dynamic changes in parasitemia, implying a correlation between peripheral densities and total parasite biomass. For the purpose of determining disease severity, total parasite biomass per kg was calculated from the predicted parasite density by the equation 70,000 × 1.09 × predicted parasite density in parasites/µL, as has been determined previously for uncomplicated malaria cases in the Gambian dataset^37^. The source code for the model and examples of its use are presented as **Supplementary Code File**.

*Deterministic relationships between real and latent variables.* The range of values of *m* and ln *P_c_* in a simulated population of 2000 patients were determined and each divided into 50 equally spaced increments in order to generate 2500 possible combinations of *m* and ln *P_c_* for which all model outcomes were determined. For the purpose of this analysis, the time-dependent adaptive immune response parameters (which comprise *S_m_*) were set for all subjects at their respective population median values. The model of Dietz *et al*. makes use of discrete 2 day time intervals^4^, corresponding to the duration of the intraerythrocytic cycle in a highly synchronized infection. However, naturally acquired infections are rarely this synchronous^38^ and the time since infection of our Gambian patients is an unknown continuous variable. In order to cope with this we assumed that the relationship between predicted outcome variables (parasite biomass, duration of illness and TNF concentration) and explanatory variables (*m* and *P_c_*) could be approximated by smoothed GAM^44^. We used the GAM to estimate values of *m, P_c_* and parasite growth inhibition (PGI, 1-*S_c_*) in the Gambian children, based on their known total parasite biomass, duration of symptoms and TNF concentration.

### RNA-sequencing and data analysis

Total RNA was extracted using the PAXgene Blood RNA kit (BD) and quality assessed using the Agilent 6000 RNA Nano kit and Bioanalyzer. Library preparation and sequencing were performed by Exeter University sequencing service. Libraries were prepared from 1μg of total RNA using the ScriptSeq v2 RNA-seq library preparation kit (Illumina) with additional steps to remove ribosmal RNA (rRNA) and globin messenger RNA (mRNA) using Globin-Zero Gold kit (Epicentre). Strand-specific libraries were sequenced using the 2×100 bp protocol with an Illumina HiSeq 2500. Samples were randomized for order of library preparation and then randomly allocated to sequencing lanes in a block design to ensure a balance of SM and UM samples (5–6 samples per lane) to eliminate batch effects. The reference genomes hg38 (http://genome.ucsc.edu/) and *P. falciparum* reference genome release 24 (http://plasmodb.org/) were used for human and parasite respectively. Human gene annotation was obtained from GENCODE (release 22) (http://gencodegenes.org/releases/) and *P. falciparum* gene annotation from PlasmoDB (release 24) (http://plasmodb.org). RNA-seq data was mapped to the combined genomic index containing both human and *P. falciparum* genomes using splice-aware STAR aligner, allowing up to 8 mismatches for each paired-end read^45^. Reads were then extracted from the output BAM file to separate parasite-mapped reads from human-mapped reads. Reads mapping to both genomes were counted for each sample and removed. BAM files were also sorted, read groups replaced with a single new read group and all reads assigned to it, and indexed to run RNASeQC, a tool for computing quality control metrics for RNA-seq data^46^. Only exclusively human-mapped reads were used for further analysis. HTSeq-count was used to count the reads mapped to exons with the parameter “-m union”^47^. With the R package edgeR, raw read counts were normalized using a trimmed mean of M-values (TMM), which takes into account the library size and the RNA composition of the input data^48^. To account for inter-individual variation in the proportions of different types of blood leukocyte, deconvolution analysis was performed using CellCODE^49^. Leukocyte expression signatures were taken from the in-built Immune Response In Silico (IRIS^50^) dataset for: neutrophil, monocyte, CD4+ T-cell, CD8+ T-cell, and B-cell. Fragments Per Kilobase of transcript per Million mapped reads (FPKM) values were calculated from human RNA-seq data and log-transformed to simulate a microarray data set. Surrogate proportion variables for each sample were calculated for each of the cell types.

The association of gene expression with *m* and PGI was determined using a generalized linear model approach in edgeR, allowing adjustment for leukocyte SPVs. False discovery rate (FDR) was computed using the Benjamini-Hochberg approach and FDR below 0.05 was considered to be significant. Gene ontology (GO) terms were obtained from Bioconductor package “org.Hs.eg.db”. Fisher’s exact test was used to identify significantly over-represented GO terms from gene lists. The background gene sets consisted of all expressed genes detected in the data set. Enrichment analysis for biological process terms was carried out using the “goana()” function in edgeR. REVIGO was used to identify the most significant non-redundant GO terms^51^. Using groups of genes significantly positively or negatively correlated with PGI, Ingenuity Pathway Analysis (Qiagen) was used to identify networks of genes functionally linked by regulators, interactions or downstream effects, which were visualized as radial plots centered around the most connected network member.

### Parasite culture, growth and invasion assays

*P. falciparum* 3D7 strain was used in continuous culture for all of the experiments unless otherwise stated. Asexual blood stage parasites were cultured in human blood group A red cells, obtained from the National Blood Service, at 1–5% hematocrit, 37 °C, 5% CO_2_ and low oxygen (1% or 5%) as described previously^52,53^. Growth medium was RPMI-1640 (without L-glutamine, with HEPES) (Sigma) supplemented with 5 g/L Albumax II (Invitrogen), 147 µM hypoxanthine, 2 mM L-glutamine, and 10 mM D-glucose. Parasite developmental stage synchronization was performed using 5% D-sorbitol to obtain ring stage parasites or Percoll gradients for schizont stage enrichment^53,54^. For growth assays, schizonts were mixed at <1% parasitemia with uninfected erythrocytes at 2% final hematocrit. Cathepsin G (Abcam) or recombinant active MMP9 [Enzo] were added for 72 hour incubation to allow two replication cycles. Growth under each condition was calculated relative to the average growth in untreated samples. Invasion assays were performed by adding parasites synchronised at the schizont stage to target erythrocytes and incubating for 24 hours. Cathepsin G and MMP9 were either pre-incubated with the target cells overnight followed by four washes with RPMI to completely remove them, or they were added directly to the culture of schizonts with target erythrocytes for 24 hours. The same protocol was followed for other *P. falciparum* strains except Dd2, for which magnetic purification was used to purify schizonts^55^.

### Flow cytometry for parasitemia and erythrocyte surface receptor expression

Flow cytometry was performed using a BD LSR Fortessa machine and analysis was conducted using FlowJo v10 (TreeStar Inc.), and gating strategies are show in Extended Data Figure 4. To assess parasitemia, 1µl of sample at 50% hematocrit was stained with Hoechst 33342 (Sigma) and dihydroethidium (Sigma) and then fixed with 2% paraformaldehyde (PFA) before flow cytometry as previously described^56^. Erythrocyte surface receptor expression was assessed by median fluorescence intensity of erythrocytes labelled with monoclonal antibodies or by comparison with isotype control antibodies (Extended Data Table 6). Briefly, erythrocytes were washed twice before resuspending at 50% haematocrit, of which 1–2µl was stained in 100µl of antibody cocktail in FACS buffer (2% fetal bovine serum, 0.01% sodium azide in PBS) for 30 minutes in the dark on ice. Samples were washed twice in FACS buffer and then fixed in 300µl FACS buffer with 2% paraformaldehyde. Surface receptor loss was calculated from the difference between the treated and untreated sample median fluorescent intensities after the isotype control antibody fluorescence had been subtracted.

### Whole blood stimulation and Cathepsin G and MMP9 ELISA

Whole blood was collected from 8 healthy adult donors and plated at 25% hematocrit, and incubated overnight with or without 1µM PMA (Sigma). Supernatant was collected to perform Cathepsin G (CTSG ELISA Kit-Human, Aviva Systems Biology) and MMP9 (Legend Max Human MMP-9, Biolegend) ELISAs, and erythrocytes were collected for surface staining. The same ELISA kits were used to measure cathepsin G and MMP9 in acute (day 0) plasma samples from children with malaria.

### Hepcidin and Complement Factor H ELISA

The hepcidin concentration was measured 28 days after infection in plasma from subjects who had not received blood transfusion using the Hepcidin-25 bioactive ELISA kit (DRG) according to the manufacturer’s instructions, with duplicate measurements when sufficient plasma was available. Complement Factor H assays were performed using an in-house ELISA as described^57^.

### Data availability

Estimates of parameters determining within-host dynamics in malariatherapy dataset were obtained from reference 4, whose corresponding author may be contacted at. Data from the Gambian subjects are presented as Supplementary Dataset. RNA-seq data have been deposited in the ArrayExpress database at EMBL-EBI (www.ebi.ac.uk/arrayexpress) under accession number E-MTAB-6413.

### Code availability

The source code for the mathematical model, and examples of its use, are presented as Supplementary material.

## Supplementary Materials

**Supplementary Fig 1.**
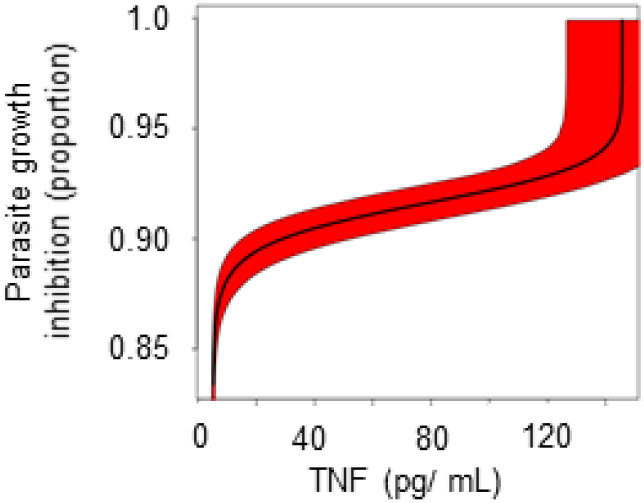
Fit of TNF concentration to parasite growth inhibition. Parameter estimation during model fitting suggested that TNF would increase above the limit of detection at a relatively late stage in the host response, when it would augment parasite killing (approximate 95% confidence interval in red)

**Supplementary Dataset. Data from the Gambian children with malaria.**

**Supplementary Code Zip File. Containing README file, R library file for the mathematical model, R Example file for the mathematical model.**

## Acknowledgments

We are grateful to Klaus Dietz for providing the original data and parameter estimates from malariatherapy patients and his model; to subjects who donated samples; to staff at MRC Gambia Unit for collection of samples; and to the St. Mary’s NHLI FACS core facility and Yanping Guo for support and instrumentation. **Funding**: This work was supported by the Medical Research Council (MRC) UK via core funding to the malaria research programme at the MRC Unit, The Gambia, and MRC Clinical Research Fellowships (G0701427 and MR/L006529/1) to A.J.C.; by a Wellcome Trust Value In People Award to A.J.C; and by European Union’s seventh Framework program under EC-GA no. 279185 (EUCLIDS; www.euclids-project.eu). **Author contributions**: A.J.C., A.G., A.E.vB., F.F., D.J.C., D.N., and M.W. collected the data used in the study; A.J.C., E.M.R., M.T.B., M.W. and D.J.C designed the study; U.D’A. and D.N. provided access to facilities and samples; A.J.C. and M.T.B. developed the mathematical model; A.J.C., A.G., M.T.B., H.J.L., F.F., and A.E.vB. analysed the data; D.W., T.W.K, D.N., U.D’A.,E.M.R., M.L., L.J.C., D.J.C., and A.J.C supervised aspects of the project; all authors contributed to interpretation of the results and drafting the manuscript. **Competing interests**: The authors declare that they have no competing financial interests.

## References

1 Plotkin, S. A. Complex correlates of protection after vaccination. Clin Infect Dis 56, 1458–1465 (2013).

2 Zumla, A. et al. Host-directed therapies for infectious diseases: current status, recent progress, and future prospects. Lancet Infect Dis 16, e47–63 (2016).

3 Cunnington, A. J. The importance of pathogen load. PLoS Pathog 11, e1004563, doi:10.1371/journal.ppat.1004563 (2015).

4 Dietz, K., Raddatz, G. & Molineaux, L. Mathematical model of the first wave of Plasmodium falciparum asexual parasitemia in non-immune and vaccinated individuals. Am J Trop Med Hyg 75, 46–55 (2006).

5 Kumaratilake, L. M., Ferrante, A. & Rzepczyk, C. M. Tumor necrosis factor enhances neutrophil-mediated killing of Plasmodium falciparum. Infect Immun 58, 788–793 (1990).

6 Kumaratilake, L. M. et al. A synthetic tumor necrosis factor-alpha agonist peptide enhances human polymorphonuclear leukocyte-mediated killing of Plasmodium falciparum in vitro and suppresses Plasmodium chabaudi infection in mice. J Clin Invest 95, 2315–2323, doi:10.1172/JCI117923 (1995).

7 Smith, T. et al. Relationships between Plasmodium falciparum infection and morbidity in a highly endemic area. Parasitology 109 (Pt 5), 539–549 (1994).

8 White, N. J. et al. Malaria. Lancet 383, 723–735 (2014).

9 Crompton, P. D. et al. Malaria immunity in man and mosquito: insights into unsolved mysteries of a deadly infectious disease. Annu Rev Immunol 32, 157–187 (2014).

10 Ceesay, S. J. et al. Continued decline of malaria in The Gambia with implications for elimination. PLoS One 5, e12242, doi:10.1371/journal.pone.0012242 (2010).

11 Okebe, J. et al. School-based countrywide seroprevalence survey reveals spatial heterogeneity in malaria transmission in the Gambia. PLoS One 9, e110926, doi:10.1371/journal.pone.0110926 (2014).

12 Arthur, J. S. & Ley, S. C. Mitogen-activated protein kinases in innate immunity. Nat Rev Immunol 13, 679–692, doi:10.1038/nri3495 (2013).

13 Manning, B. D. & Toker, A. AKT/PKB Signaling: Navigating the Network. Cell 169, 381–405 (2017).

14 Spaulding, E. et al. STING-Licensed Macrophages Prime Type I IFN Production by Plasmacytoid Dendritic Cells in the Bone Marrow during Severe Plasmodium yoelii Malaria. PLoS Pathog 12, e1005975, doi:10.1371/journal.ppat.1005975 (2016).

15 Zander, R. A. et al. Type I Interferons Induce T Regulatory 1 Responses and Restrict Humoral Immunity during Experimental Malaria. PLoS Pathog 12, e1005945, doi:10.1371/journal.ppat.1005945 (2016).

16 Haque, A. et al. Type I IFN signaling in CD8-DCs impairs Th1-dependent malaria immunity. J Clin Invest 124, 2483–2496 (2014).

17 Haque, A. et al. Type I interferons suppress CD4(+) T-cell-dependent parasite control during blood-stage Plasmodium infection. Eur J Immunol 41, 2688–2698 (2011).

18 Edwards, C. L. et al. Spatiotemporal requirements for IRF7 in mediating type I IFN-dependent susceptibility to blood-stage Plasmodium infection. Eur J Immunol 45, 130–141 (2015).

19 Montes de Oca, M. et al. Type I Interferons Regulate Immune Responses in Humans with Blood-Stage Plasmodium falciparum Infection. Cell reports 17, 399–412 (2016).

20 Ioannidis, L. J. et al. Monocyte-and Neutrophil-Derived CXCL10 Impairs Efficient Control of Blood-Stage Malaria Infection and Promotes Severe Disease. J Immunol 196, 1227–1238 (2016).

21 Cowland, J. B. & Borregaard, N. Granulopoiesis and granules of human neutrophils. Immunol Rev 273, 11–28, doi:10.1111/imr.12440 (2016).

22 Weksler, B. B., Jaffe, E. A., Brower, M. S. & Cole, O. F. Human leukocyte cathepsin G and elastase specifically suppress thrombin-induced prostacyclin production in human endothelial cells. Blood 74, 1627–1634 (1989).

23 Binks, R. H. & Conway, D. J. The major allelic dimorphisms in four Plasmodium falciparum merozoite proteins are not associated with alternative pathways of erythrocyte invasion. Mol Biochem Parasitol 103, 123–127 (1999).

24 Satchwell, T. J. Erythrocyte invasion receptors for Plasmodium falciparum: new and old. Transfusion medicine 26, 77–88 (2016).

25 Malaria Genomic Epidemiology, N., Band, G., Rockett, K. A., Spencer, C. C. & Kwiatkowski, D. P. A novel locus of resistance to severe malaria in a region of ancient balancing selection. Nature 526, 253–257 (2015).

26 Leffler, E. M. et al. Resistance to malaria through structural variation of red blood cell invasion receptors. Science, eaam6393, doi:10.1126/science.aam6393 (2017).

27 Crosnier, C. et al. Basigin is a receptor essential for erythrocyte invasion by Plasmodium falciparum. Nature 480, 534–537 (2011).

28 Pasvol, G., Wainscoat, J. S. & Weatherall, D. J. Erythrocytes deficiency in glycophorin resist invasion by the malarial parasite Plasmodium falciparum. Nature 297, 64–66 (1982).

29 Clark, M. A. et al. Host iron status and iron supplementation mediate susceptibility to erythrocytic stage Plasmodium falciparum. Nature communications 5, 4446, doi:10.1038/ncomms5446 (2014).

30 Kennedy, A. T. et al. Recruitment of Factor H as a Novel Complement Evasion Strategy for Blood-Stage Plasmodium falciparum Infection. J Immunol 196, 1239–1248 (2016).

31 Rosa, T. F. et al. The Plasmodium falciparum blood stages acquire factor H family proteins to evade destruction by human complement. Cell Microbiol 18, 573–590 (2016).

32 Gwamaka, M. et al. Iron deficiency protects against severe Plasmodium falciparum malaria and death in young children. Clin Infect Dis 54, 1137–1144 (2012).

33 Pasricha, S. R. et al. Expression of the iron hormone hepcidin distinguishes different types of anemia in African children. Sci Transl Med 6, 235re233, doi:10.1126/scitranslmed.3008249 (2014).

34 Parente, R., Clark, S. J., Inforzato, A. & Day, A. J. Complement factor H in host defense and immune evasion. Cell Molecular Life Sci 74, 1605–1624 (2017).

35 Oon, S., Wilson, N. J. & Wicks, I. Targeted therapeutics in SLE: emerging strategies to modulate the interferon pathway. Clin Transl Immunol 5, e79, doi:10.1038/cti.2016.26 (2016).

36 Pease, J. E. Designing small molecule CXCR3 antagonists. Expert Opin Drug Discov 12, 159–168 (2017).

37 Cunnington, A. J., Bretscher, M. T., Nogaro, S. I., Riley, E. M. & Walther, M. Comparison of parasite sequestration in uncomplicated and severe childhood Plasmodium falciparum malaria. J Infect 67, 220–230 (2013).

38 Dondorp, A. M. et al. Estimation of the total parasite biomass in acute falciparum malaria from plasma PfHRP2. PLoS medicine 2, e204, doi:10.1371/journal.pmed.0020204 (2005).

39 Griffin, J. T. et al. Gradual acquisition of immunity to severe malaria with increasing exposure. Proc Biol Sci 282, doi:10.1098/rspb.2014.2657 (2015).

40 Walther, M. et al. HMOX1 gene promoter alleles and high HO-1 levels are associated with severe malaria in Gambian children. PLoS Pathog 8, e1002579, doi: 10.1371/journal.ppat.1002579 (2012).

41 Walther, M. et al. Distinct roles for FOXP3 and FOXP3 CD4 T cells in regulating cellular immunity to uncomplicated and severe Plasmodium falciparum malaria. PLoS Pathog 5, e1000364, doi:10.1371/journal.ppat.1000364 (2009).

42 Marsh, K. et al. Indicators of life-threatening malaria in African children. N Engl J Med 332, 1399–1404 (1995).

43 R Development Core Team. R: A language and environment for statistical computing. R Foundation for Statistical Computing, <URL http://www.R–project.org/> (2014).

44 Wood, S. N. Fast stable restricted maximum likelihood and marginal likelihood estimation of semiparametric generalized linear models. J R Stat Soc B 73, 3–36 (2011).

45 Dobin, A. et al. STAR: ultrafast universal RNA-seq aligner. Bioinformatics 29, 15–21 (2013).

46 DeLuca, D. S. et al. RNA-SeQC: RNA-seq metrics for quality control and process optimization. Bioinformatics 28, 1530–1532 (2012).

47 Anders, S., Pyl, P. T. & Huber, W. HTSeq--a Python framework to work with high-throughput sequencing data. Bioinformatics 31, 166–169 (2015).

48 Robinson, M. D., McCarthy, D. J. & Smyth, G. K. edgeR: a Bioconductor package for differential expression analysis of digital gene expression data. Bioinformatics 26, 139–140 (2010).

49 Chikina, M., Zaslavsky, E. & Sealfon, S. C. CellCODE: a robust latent variable approach to differential expression analysis for heterogeneous cell populations. Bioinformatics 31, 1584–1591 (2015).

50 Abbas, A. R. et al. Immune response in silico (IRIS): immune-specific genes identified from a compendium of microarray expression data. Genes Immun 6, 319–331 (2005).

51 Supek, F., Bosnjak, M., Skunca, N. & Smuc, T. REVIGO summarizes and visualizes long lists of gene ontology terms. PLoS One 6, e21800, doi:10.1371/journal.pone.0021800 (2011).

52 Trager, W. & Jensen, J. B. Human malaria parasites in continuous culture. Science 193, 673–675 (1976).

53 Maier, A. G. & Rug, M. In vitro culturing Plasmodium falciparum erythrocytic stages. Methods Mol Biol 923, 3–15 (2013).

54 Dluzewski, A. R., Ling, I. T., Rangachari, K., Bates, P. A. & Wilson, R. J. A simple method for isolating viable mature parasites of Plasmodium falciparum from cultures. Trans R Soc Trop Med Hyg 78, 622–624 (1984).

55 Ribaut, C. et al. Concentration and purification by magnetic separation of the erythrocytic stages of all human Plasmodium species. Malar J 7, 45, doi:10.1186/1475-2875-7-45 (2008).

56 Malleret, B. et al. A rapid and robust tri-color flow cytometry assay for monitoring malaria parasite development. Sci Rep 1, 118, doi:10.1038/srep00118 (2011).

57 Pouw, R. B. et al. Complement Factor H-Related Protein 3 Serum Levels Are Low Compared to Factor H and Mainly Determined by Gene Copy Number Variation in CFHR3. PLoS One 11, e0152164, doi:10.1371/journal.pone.0152164 (2016).

